# Class IV plant U-box proteins function redundantly to optimize protein accumulation of receptor-like cytoplasmic kinase BIK1

**DOI:** 10.1101/2025.02.01.635694

**Authors:** Ruoqi Dou, Virginia N. Miguel, Lauren E. Grubb, Mansuba Rana, Brandon Saltzman, Jacqueline Monaghan

## Abstract

In *Arabidopsis thaliana*, the receptor-like cytoplasmic kinase BOTRYTIS INDUCED KINASE 1 (BIK1) is a direct substrate of multiple transmembrane immune receptor kinases and plays a crucial role in immune signal transduction. Inactive BIK1 is poly-ubiquitinated and degraded by the 26S proteasome, which is thought to optimize BIK1 levels in naive cells and may protect against inappropriately high immune responses. Here, we provide biochemical and genetic evidence that supports redundant roles between related Plant U-Box (PUB) proteins PUB22, PUB23, PUB24, PUB25, and PUB26 in BIK1 turnover.

## Main text

In *Arabidopsis thaliana*, the receptor-like cytoplasmic kinase BOTRYTIS INDUCED KINASE 1 (BIK1) is a direct substrate of multiple transmembrane immune receptor kinases including FLAGELLIN SENSING 2 (FLS2) and ELONGATION FACTOR TU RECEPTOR (EFR) (DeFalco and Zipfel, 2021). BIK1 plays a crucial role in immune signal transduction by phosphorylating multiple substrates, including the NADPH oxidase RESPIRATORY OXIDASE HOMOLOG D (RBOHD) which catalyzes the production of reactive oxygen species (ROS) in the apoplast (Gonçalves Dias et al., 2022). BIK1 accumulation is tightly controlled via both proteasomal (Monaghan et al., 2014; Wang et al., 2018; Yu et al., 2022) and endosomal mechanisms (Ma et al., 2020). Inactive BIK1 is poly-ubiquitinated and degraded by the 26S proteasome, which is thought to optimize BIK1 levels in naive cells and may protect against inappropriately high immune responses. Ubiquitination is mediated by multiple families of E3 ligases, including several classes of plant U-box (PUB) proteins that pair with E2 conjugating enzymes to transfer ubiquitin to target proteins (Trenner et al., 2022). Several class IV PUBs (**Figure 1A**) negatively regulate immune signaling, including paralogs PUB20 and PUB21 (Yi et al., 2024), PUB22, PUB23, and PUB24 (Trujillo et al., 2008), and PUB25 and PUB26 (Wang et al., 2018). PUB25 and PUB26 poly-ubiquitinate BIK1 and are directly phosphorylated by the calcium-dependent protein kinase CPK28 (Wang et al., 2018). In addition, over-expression of *PUB22* or *PUB23* alters BIK1 stability in Arabidopsis protoplasts (Wang et al., 2018). Here, we provide biochemical and genetic evidence that supports redundant roles between PUB22, PUB23, PUB24, PUB25, and PUB26 in BIK1 turnover.

**Figure 1.**
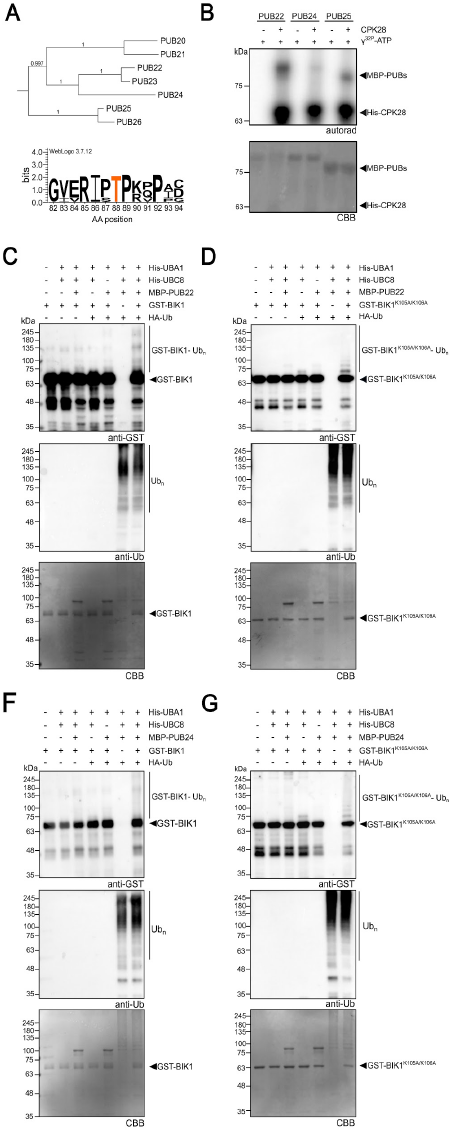
PUB22 and PUB24 are phosphorylated by CPK28 and ubiquitinate BIK1 *in vitro*. **(A)** Phylogenetic tree of a subgroup of class IV PUBs using sequences obtained from The Arabidopsis Information Resource (TAIR) according to (Trenner et al., 2022). Protein sequences were aligned with MAFFT v7.490, and a maximum-likelihood tree was constructed using IQ-TREE v2.3.4 with 1,000 bootstraps. The final tree was visualized and annotated in iTOL v6. The WebLogo illustrates conserved positions from a multiple protein sequence alignment of PUB22, PUB23, PUB24, PUB25, and PUB26, with the conserved Thr (T) highlighted in orange. Bit scores are indicated on the y-axis, where a score of 4 indicates 100% conservation, and amino acid positions are indicated on the x-axis according to their position in PUB22. Analysis and figure generated by VNM. **(B)** *In vitro* kinase assays with His-CPK28 as the kinase and MBP-tagged PUB22, PUB24 and PUB25 as substrates. Phosphorylation was detected via incorporation of ^32^P as visualized by autoradiography (autorad). Protein loading was verified by post-staining the gel with Coomassie Brilliant Blue (CBB); note that CPK28 is not visible due to the small amount loaded. Assays were performed over six times by RD and VNM with consistent results. **(C-F)** Ubiquitination assays using His-UBC1 (E1), His-UBC8 (E2), and either MBP-PUB22 (E3) **(C-D)** or MBP-PUB24 (E3) **(E-F)**, with GST-BIK1 **(C,E)** or GST-BIK1^K105A/K106A^ **(D,F)** as substrates. Western blots were probed with anti-GST or anti-Ub antibodies; CBB staining of the membrane indicates protein loading. Assays were repeated more than three times with similar results by RD.

CPK28 phosphorylates PUB25 and PUB26 on multiple residues (Wang et al., 2018). Among these, PUB25-Thr95 and PUB26-Thr94 are in conserved positions and their phosphorylation contributes to catalytic activation (Wang et al., 2018). In addition, the mitogen-activated protein kinase MPK3 phosphorylates PUB22 on the orthologous residue Thr88 (Furlan et al., 2017), and phosphorylation on PUB25-Thr95, PUB26-Thr94, and PUB22-Thr88 is induced *in vivo* following treatment with the immunogenic elicitor flg22 (the 22-amino acid active epitope of bacterial flagellin) (Furlan et al., 2017; Wang et al., 2018). Furthermore, it was recently reported that phosphorylation at this residue is targeted by the effector protein RipAV from the vascular bacterial pathogen *Ralstonia solanacearum*, resulting in decreased accumulation of BIK1 (Rufian et al., 2025).

As this residue is conserved in PUB22, PUB23, PUB24, PUB25, and PUB26 (**Figure 1A**), we hypothesized that other members of this class may also be phosphorylated by CPK28. We therefore selected PUB22 and PUB24 to test as additional substrates of CPK28 and found that both can indeed be phosphorylated by CPK28 *in vitro* (**Figure 1B**). We next assessed whether PUB22 and PUB24 can ubiquitinate BIK1 *in vitro*. As PUB25 and PUB26 preferentially poly-ubiquitinate inactive variants of BIK1 (Wang et al., 2018), we tested both catalytically active (BIK1) and inactive (BIK1^K105A/K106A^) variants of BIK1 as substrates of PUB22 and PUB24. We found that PUB22 and PUB24 can poly-ubiquitinate both variants when paired with the E2 ubiquitin-conjugating (UBC) protein UBC8 (**Figure 1C-F**). Taken together, we conclude that multiple class IV PUBs can be phosphorylated by CPK28 and are capable of poly-ubiquitinating BIK1 *in vitro*.

Immune signaling is deregulated in *pub22 pub23 pub24* (Trujillo et al., 2008) and *pub25 pub26* (Wang et al., 2018) mutants, presenting as increased immune-triggered ROS production and resistance to pathogen infection. Whilst the enhanced immune phenotypes observed in *pub25 pub26* are less pronounced than in *pub22 pub23 pub24*, stable over-expression of *PUB25* or *PUB26* results in strongly reduced immune responses (Wang et al., 2018). To further test functional redundancy between class IV PUBs, we generated a *pub22 pub23 pub24 pub25 pub26* quintuple mutant and assessed immune responses. In wild-type plants, ROS production peaks by ∼10 minutes following immune detection and returns to baseline by ∼40 minutes. We found that the quintuple mutant presents an extremely high level of ROS production and a very slow rate of decline following exposure to the immunogenic elicitors flg22 or elf18 (the 18 amino-acid active epitope of bacterial Elongation Factor Tu) when compared to both *pub22 pub23 pub24* and *pub25 pub26* (**Figure 2A-B; Figure S1A-B**). The slower rate of decline is congruent with what has been previously described in *pub22 pub23 pub24* (Trujillo et al., 2008). We also observed a second elf18-induced ROS peak after ∼120 minutes in all genotypes, however, this occurred earlier and with a much higher amplitude in both *pub22 pub23 pub24* and *pub22 pub23 pub24 pub25 pub26* mutants (**Figure 2A**). Intriguingly, we also observed a third ROS peak following elf18 treatment in *pub22 pub23 pub24* and *pub22 pub23 pub24 pub25 pub26* mutants, which was extremely high in the quintuple mutant (**Figure 2A**). These results support functional redundancy between class IV PUBs and furthermore suggest divergent mechanisms of signal attenuation following FLS2 or EFR activation. Resistance to the virulent bacterial pathogen *Pseudomonas syringae* pathovar *tomato* DC3000 (*P.s.t*. DC3000) was also strongly enhanced in *pub22 pub23 pub24 pub25 pub26* compared to Col-0, *pub22 pub23 pub24*, or *pub25 pub26* (**Figure 2C**; **Figure S2**). Remarkably, enhanced immune responses in *pub22 pub23 pub24 pub25 pub26* do not correlate with typical complications of autoimmunity such as stunted growth (**Figure 2D**).

**Figure 2.**
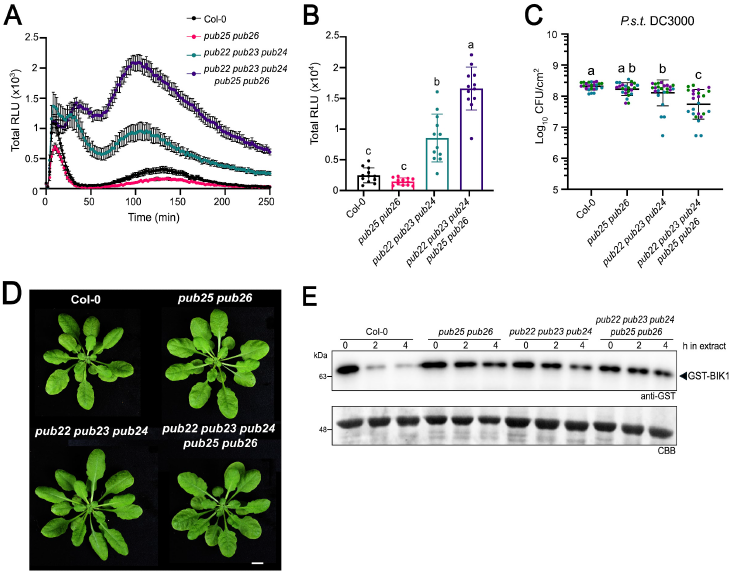
Class IV PUBs function redundantly to optimize protein accumulation of receptor-like cytoplasmic kinase BIK1. **(A-B)** ROS production measured in relative light units (RLUs) after treatment with 50 µM elf18. Values in **A** represent means +/-standard error (n=12 leaf discs) measured every 2 minutes; values in **B** represent means +/-standard deviation (n=12) of total RLU captured in 250 minutes. Lower-case letters indicate statistically significant groups determined by a one-way ANOVA followed by Tukey’s posthoc test (p-value < 0.0001). Error bars represent standard deviation. These assays were conducted more than 10 times by BS, RD, and VNM; representative data collected by VNM is shown. **(C)** Growth of virulent *Pseudomonas syringae* pv. *tomato* (*P.s.t*.) isolate DC3000 three days after syringe-infiltration using a bacterial suspension of 10^5^ cfu/mL. Data from 3 independent experimental replicates are plotted together, denoted by blue, green, and purple dots. Values are log_10_ of colony-forming units (CFU) per leaf area (cm^2^) from 4 samples per genotype (each sample contains 3 leaf discs from 3 different infected plants). Error bars represent standard deviation (n=24). Lower-case letters indicate statistically significant groups determined by a one-way ANOVA followed by Tukey’s post-hoc test (p-value < 0.0001). Data collected by VNM. **(D)** Images of 4-week-old plants grown under short-day conditions, taken by VNM. The white bar represents 1 cm. **(E)** Anti-GST western blots following the cell-free degradation of GST-BIK1 in total protein extracts from Col-0, *pub25 pub26, pub22 pub23 pub24*, and *pub22 pub23 pub24 pub25 pub26* over 0, 2, or 4 hours. Post-staining the membrane with Coomassie Brilliant Blue (CBB) indicates loading. Assays were performed over five times by RD with consistent results. LEG, MR, and JM performed genetic crosses and genotyping to isolate the *pub22 pub23 pub24 pub25 pub26* quintuple mutant as indicated in more detail in **Table S1**.

Based on our results, we hypothesized that subclass IV PUBs redundantly regulate BIK1 accumulation. A useful way to test this is via cell-free degradation assays, in which recombinant proteins are bathed in plant protein extracts and their stability is monitored over time by immunoblot. We know from previous work that recombinant BIK1 degrades rapidly in Col-0 protein extract, which can be suppressed using proteasomal inhibitors (Liang et al., 2016). We found that the rate of BIK1 degradation is slower in *pub22 pub23 pub24, pub25 pub26*, and *pub22 pub23 pub24 pub25 pub26* extracts compared to Col-0 extracts (**Figure 2E**), providing evidence that they contribute to BIK1 turnover. Overall, our results suggest that PUB22, PUB23, PUB24, PUB25, and PUB26 redundantly contribute to the attenuation and/or buffering of plant immune responses by regulating the accumulation of BIK1.

## Acknowledgements

We thank all members of the Monaghan Lab for their commitment to fostering a welcoming and collaborative research environment. We thank Marco Trujillo for sharing published materials and Saeid Mobini for managing the Queen’s University Phytotron Facility. Queen’s University is situated on the territory of the Haudenosaunee and Anishinaabek and we are grateful to live, work, and play on these lands.

## Supplemental Information

The following supplemental information is available for this manuscript:

- **Table S1**. Germplasm, reagents, clones, and primers used in this study.
- **Figure S1.** Flg22-induced ROS in class-IV PUB mutants.
- **Figure S2.** Disease symptoms in class IV PUB mutants infected with *P.s.t*. DC3000.
- **Supplemental Methods**.
- **Author Contributions**.
- **Supplemental References**.

