## Supplemental Information for "Class IV plant U-box proteins function redundantly to optimize protein accumulation of receptor-like cytoplasmic kinase BIK1"

---

### Supplemental Information

The following supplemental information is available for this manuscript:

- **Table S1**. Germplasm, reagents, clones, and primers used in this study.
- **Figure S1**. Flg22-induced ROS in class IV PUB mutants.
- **Figure S2**. Disease symptoms in class IV PUB mutants infected with *P.s.t.* DC3000.
- **Supplemental Methods**.
- **Author Contributions.**
- **Supplemental References.**

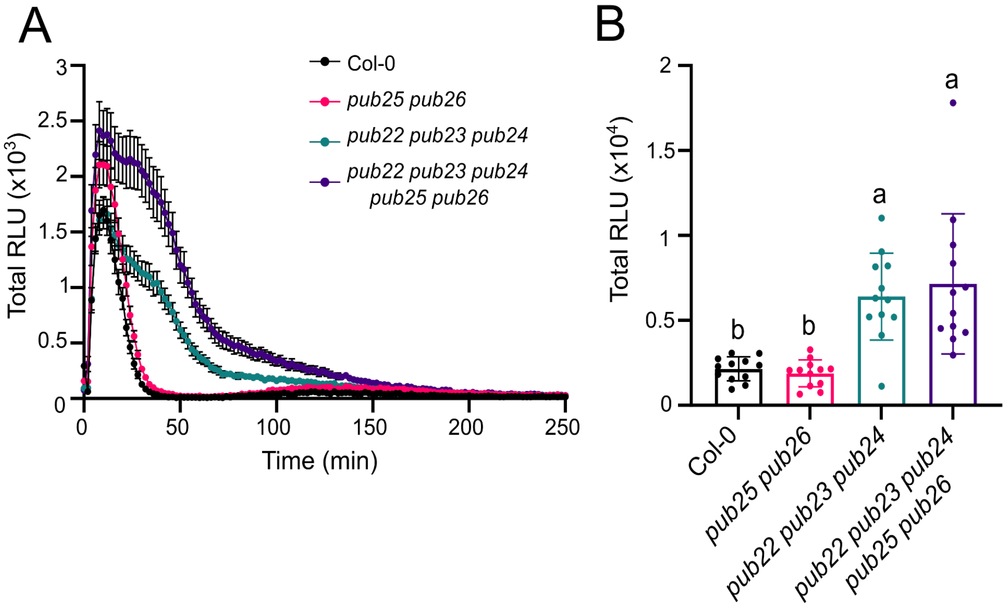


**Figure S1. Flg22-induced ROS in class IV PUB mutants.**

**(A-B)** ROS production measured in response to 50 μM flg22 over 250 minutes, visualized as relative light units (RLU). Values in **A** represent means +/- standard error (n=12 leaf discs) measured every 2 minutes; values in **B** represent means +/- standard deviation (n=12) of total RLU captured in 250 minutes. Lower-case letters indicate statistically significant groups determined by a one-way ANOVA followed by Tukey's posthoc test (p-value < 0.0001). These assays were conducted more than ten times by BS, RD, and VNM; representative data collected by VNM is shown.

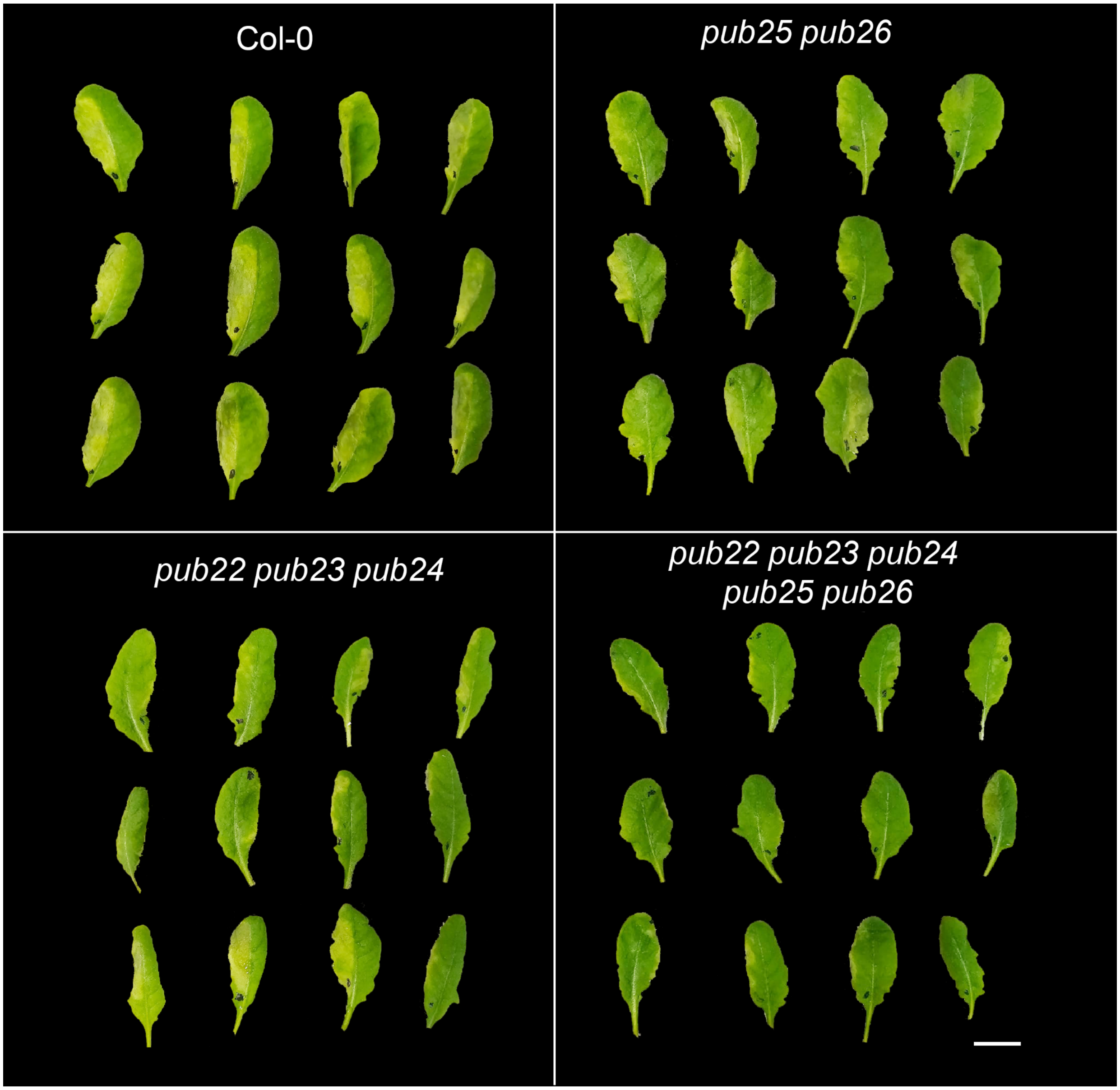


**Figure S2. Disease symptoms in class IV PUB mutants infected with *P.s.t.* DC3000.** Six-week-old plants of the indicated genotypes three days after infiltration with *Pseudomonas syringae* pv. *tomato* DC3000 (*P.s.t.* DC3000). Black dots indicate which side of the leaf was infiltrated and the white bar represents 1 cm. Photos are representative of three experiments conducted by VNM.

### Supplemental Methods

#### Plant materials, growth conditions, and physiological assays

*Arabidopsis thaliana* plants were grown in controlled growth chambers under short-day conditions (10 h light; 14 h dark) as recently described (Gonçalves Dias et al., 2024). Detailed information about all germplasm used in this study, including previously published materials and primers used for genotyping, is provided in **Table S1**. To generate the *pub22 pub23 pub24 pub25 pub26* quintuple mutant, LEG crossed homozygous *pub22 pub23 pub24* (Trujillo et al., 2008) with homozygous *pub25 pub26* (Wang et al., 2018) insertional mutants, and LEG, MR, and JM genotyped subsequent segregating populations using gene- and insert-specific primers through the $F_4$ generation until individuals were identified as homozygous for all loci. Flg22 and elf18 peptides were synthesized by EZ Biolabs (USA) and ROS production was measured by VNM, RD, and BS using a luminol-based assay as previously described (Bredow et al., 2019). Syringe-inoculation of $10^5$ colony-forming units per mL *Pseudomonas syringae* pv. *tomato* DC3000 was performed by VNM on 4- to 5-week-old soil-grown Arabidopsis plants exactly as described in (Monaghan et al., 2014). Cell-free degradation assays were performed by RD exactly as recently described (Dou et al., 2024), using tissue from 4- to 5-week-old healthy mid-layer leaves from soil-grown Arabidopsis plants.

### Recombinant protein purification and *in vitro* ubiquitination and kinase assays

All clones used in this study were previously published and are listed with citation details in **Table S1**. Protein expression was induced with isopropyl β- d-1-thiogalactopyranoside (IPTG) in *Escherichia coli* BL21 cells and purified according to standard protocols, exactly as described recently (Gonçalves Dias et al., 2024). $His_6$-tagged proteins were purified using nickel-nitrilotriacetic acid resin (HisPur™ Ni-NTA, Thermo Fisher Scientific); GST-tagged proteins were purified with pre-hydrated glutathione agarose beads (Sigma Aldrich); and MBP-tagged proteins were purified using amylose resin (New England Biolabs). *In vitro* kinase assays were performed as recently described (Gonçalves Dias et al., 2024), using 0.5 ng of kinase and 4 µg of substrate in a buffer containing 50 mM Tris-HCl (pH 8.0), 25 mM $MgCl_2$, 500 µM $CaCl_2$, 5 mM DTT, 5 µM ATP, and 2 µCi γ-[$^{32}$P]-ATP, and incubated at 30°C for 10 minutes. Reactions were terminated in 5x Laemmli Sample Buffer at 80°C for 5 minutes and proteins were separated by 6% SDS-PAGE. The resulting gels were sandwiched between two sheets of transparency film, exposed to a storage phosphor screen (Molecular Dynamics), and visualized using a Typhoon 9700 Imager (Molecular Dynamics). Gels were post-stained with Coomassie Brilliant Blue (CBB) R-250 (MP Biomedicals) and scanned. *In vitro* ubiquitination assays and immunoblots were performed exactly as recently described (Dou et al., 2024), using 2 µg of recombinant MBP-PUB22 or MBP-PUB24, 4 µg of GST-BIK1 or GST-BIK1$^{K105A/K106A}$, and 3.75 µg of HA-tagged ubiquitin (R&D Systems) in a 60 µL buffer containing 25 mM Tris-HCl (pH 7.4), 5 mM $MgCl_2$, 25 mM KCl, 0.33 mM DTT, 1.5 mM ATP, 200 ng E1 (UBA1), and 450 ng E2 (UBC8) for 180 min. Reactions were terminated by adding 5×

Laemmli Sample Buffer, followed by heating at 80°C for 5 minutes. A 20 µL aliquot of each reaction mixture was subjected to immunoblot analysis. Proteins were separated using 10% SDS-PAGE, transferred onto polyvinylidene difluoride (PVDF) membranes (Bio-Rad), and blocked with 5% nonfat milk in Tris-buffered saline containing 0.1% Tween-20. Membranes were probed with anti-Ub (P4D1, Cell Signaling Technology) and anti-GST (Sigma-Aldrich) antibodies, incubated with ECL Clarity Substrate (Bio-Rad), and visualized using the ChemiDoc Touch Imaging System (Bio-Rad). As a loading control, membranes were subsequently stained with Coomassie Brilliant Blue (CBB) R-250 (MP Biomedicals). See **Table S1** for more information on reagents.

### Data analysis and statistics

Data visualization and statistics were performed using GraphPad Prism v10. See figure captions for more information.

### Phylogenetics

Protein sequences of class IV PUBs were selected according to (Trenner et al., 2022) and aligned using MAFFT v7.490 (Katoh and Standley, 2013). A maximum-likelihood phylogenetic tree was then constructed with IQ-TREE v2.3.4 (Minh et al., 2020) using 1,000 bootstraps and visualized using iTOL v6 (Letunic and Bork, 2021). All sequences were obtained from The Arabidopsis Information Resource (TAIR).

### Author contributions

**RD, VNM** - methodology, investigation, writing; **LEG, MR** - methodology; **BS** - investigation; **JM** - methodology, investigation, writing, supervision, funding acquisition. All authors contributed to editing the final version of the manuscript.
